# Applications of Wedderburn’s Theorem in Modelling Non-Commutative Biological and Evolutionary Systems

**DOI:** 10.1101/2025.04.03.647006

**Authors:** Arturo Tozzi

## Abstract

A wide range of biological and evolutionary processes is determined not merely by the occurrence of specific events, but by the exact order in which those events unfold. Gene regulation, developmental pathways, metabolic cascades and genetic evolution often display non-commutative behaviour, in which reversing the sequence of events results in functionally distinct outcomes. Conventional modelling approaches often fail to account for such directionality and sequence dependence, thereby limiting their capacity to capture the complexity of regulatory logic. We represent ordered sequences of biological operations—such as transcription factor binding or mutational trajectories—as elements of a non-commutative algebra designed to encode the functional logic of systems where the order of events is critically determinant. Subsequently, we apply Wedderburn’s Theorem to decompose the algebra into a direct sum of matrix modules, each capturing an irreducible and functionally distinct component of the underlying sequence-dependent system. We provide examples from gene expression regulation and evolutionary dynamics, focusing on scenarios where trait development is determined by the specific order of underlying molecular or mutational events. Our results demonstrate that the algebraic framework effectively maps intricate biological processes onto smaller, linear components, facilitating clearer interpretation and analysis. Simplifying non-commutative biological systems into interpretable submodules, Wedderburn-style decomposition may clarify gene regulatory logic, capture behavioural outcomes, reduce computational burden and uncover pathway redundancies and structural symmetries. Overall, by unifying diverse biological processes within a coherent algebraic structure, our method may improve the tractability of complex, order-dependent systems.

## INTRODUCTION

Recent advances in systems biology and evolutionary theory have highlighted the importance of order-dependence and context sensitivity in biological processes. Gene regulation, signal transduction, developmental cascades and adaptive evolution exhibit behaviors that resist full explanation through classical commutative models. For instance, the outcomes of regulatory or mutational sequences often depend not only on which events occur but on the precise order in which they unfold—an inherently non-commutative property (Buenrostro et al., 2018; Pountain et al., 2024). Traditional approaches such as Boolean networks, differential equations or stochastic models obscure or simplify these order effects, limiting their capacity to capture the deeper algebraic structures underlying biological causality (Chakrabarty et al., 2016; Blomberg et al., 2020; Pušnik et al., 2022; Plaugher and Murrugarra 2023). While several abstract formalisms, including category theory, non-commutative probability and rule-based modeling have been explored to address such complexities, a systematic algebraic framework assessing the modular and irreducible components of these processes remains underdeveloped (Wilson-Kanamori et al., 2015; Bustos et al., 2018; Marcot 2021). We introduce an application of the Wedderburn–Artin theorem, a foundational result in the structure theory of algebras, to decompose non-commutative biological and evolutionary systems into direct sums of matrix algebras (Lam 2001; Brešar, 2024.). This approach may allow complex, sequence-sensitive pathways to be analyzed through the lens of finite-dimensional semisimple algebras. This approach may establish a concrete algebraic foundation for modeling non-commutative biological orderings in a decomposable and interpretable form.

We propose a framework in which biological or evolutionary operations—such as binding events, mutational steps or regulatory transitions—are encoded as elements of a non-commutative associative algebra. By treating these operations as algebraic generators and constructing the corresponding finite-dimensional algebra, we aim to capture both the sequence-dependence and the compositional rules of the system. Applying Wedderburn’s theorem to these algebras yields a decomposition into matrix modules that each act irreducibly on distinct subspaces of the system’s state space (Behboodi et al., 2018; Ma et al., 2022). This decomposition enables a modular interpretation of complex pathways, revealing redundancy, symmetry and irreducibility across diverse biological sequences. We expect our experimental models to demonstrate that even relatively intricate biological systems—such as epistatic mutation chains or order-dependent regulatory switches (Wan et al., 2018; Alfaro-Murillo and Townsend, 2023)—can be mapped onto manageable algebraic structures where behavioral modules are linearly represented. These outcomes may improve the accessibility of biological models to formal analysis and the detection of functional equivalence among different sequences.

We will proceed as follows: first, we outline the algebraic formalism and its biological interpretation; second, we construct illustrative systems and apply Wedderburn’s theorem; third, we examine the resulting matrix decompositions and their relevance; finally, we discuss limitations and interpret the broader implications.

## MATERIALS AND METHODS

### Algebraic construction of biological systems

The algebraic modeling in this study is grounded in the construction of finite-dimensional associative algebras over a base field 𝔽 taken throughout as ℝ unless otherwise stated (Alder and Volker Strassen, 1981). **Figure 1** provides a flowchart of the steps described in this section. The modeling process begins by identifying a finite set of biological or evolutionary operations, denoted 𝒢 ={*g*_1_, *g*_2,…,_ *g*_*n*_}where each generator *g*_*i*_ corresponds to an ordered event in a biological pathway, such as a mutation, a transcription factor binding or a signal transduction step. These generators form the basis of a free algebra 𝔽 ⟨𝒢 ⟩which consists of all finite linear combinations of words formed from the *g*_*i*_ under non-commutative multiplication (Kleyn 2018). The product of two words 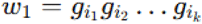 and 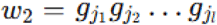 is defined by concatenation: 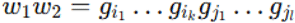. We then impose relations *R* ⊆ 𝔽 ⟨𝒢 ⟩which encode the biological logic and constraints, such as idempotency 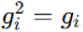, interaction rules *g*_*i*_ *g*_*j*_ = 0 if *g*_*i*_ negates *g*_*i*_ or commutativity *g*_*i*_ *g*_*j*_ = *g*_*i*_ *g*_*j*_ in the rare cases where ordering is irrelevant. The quotient algebra 𝒜= 𝔽 ⟨ 𝒢 ⟩/ ⟨ *R* ⟩ serves as the structural representation of the biological system. The definition of the quotient algebra is essential as it allows for the reduction of the free algebra into a finite-dimensional associative algebra preserving the semantic content of the biological constraints (Bremner 2010). This construction defines the primary algebraic structure upon which Wedderburn decomposition is applied, serving as the basis for all subsequent structural analysis (Brochero Martínez et al., 2022).

**Figure 1.**
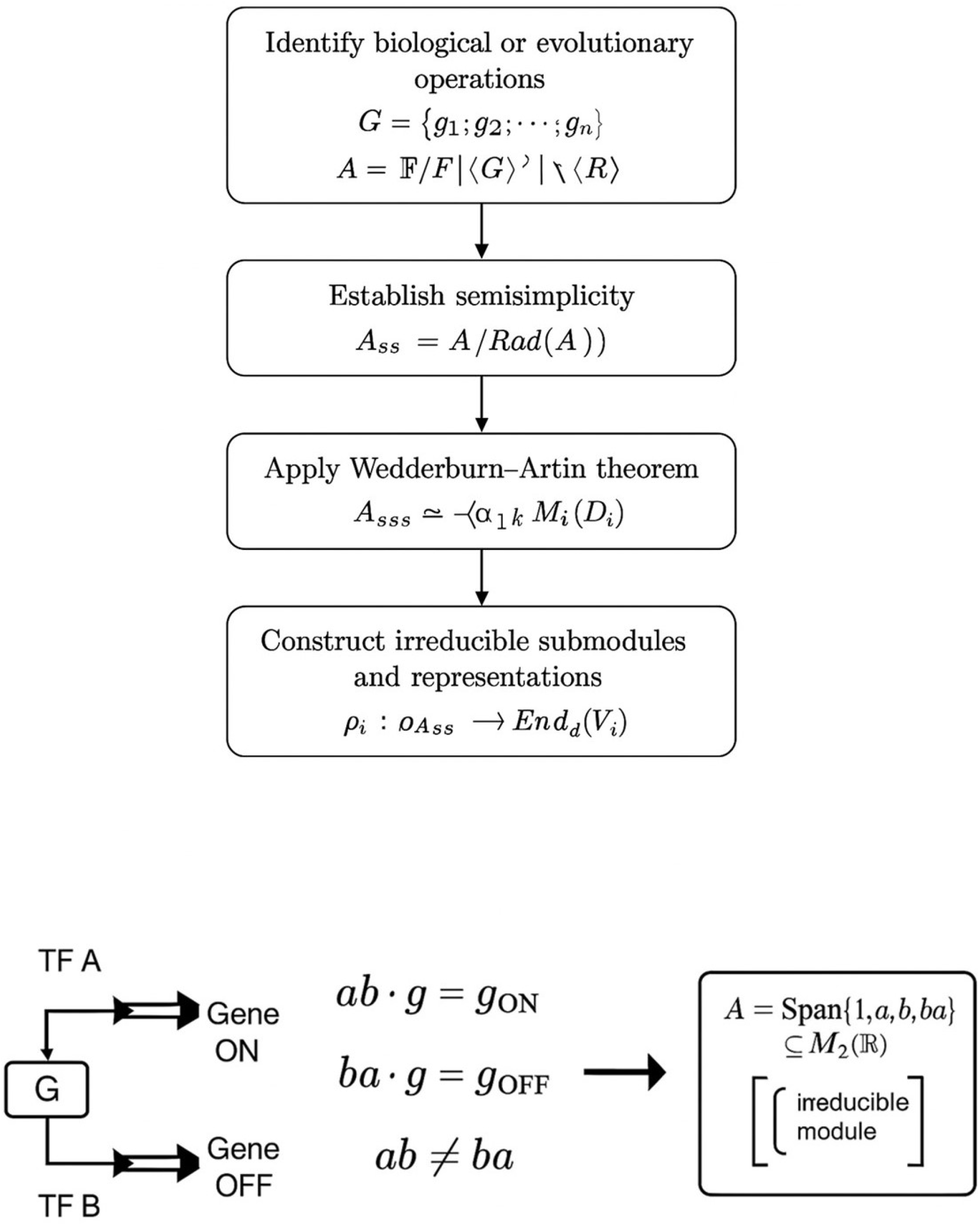
Wedderburn-style decomposition. **Upper figure**. Flowchart detailing the steps described in the Materials and Methods. **Lower figure**. A simplified gene regulation scenario illustrating non-commutative biological behavior. The sequence of transcription factor binding events (A before B vs. B before A) leads to different gene expression states. Representing these operations algebraically as non-commutative elements a and b, the resulting algebra 𝒜 admits a matrix representation and, under suitable conditions, decomposes via Wedderburn’s Theorem. This reveals underlying structural properties of the regulatory system.

**Figure 2.**
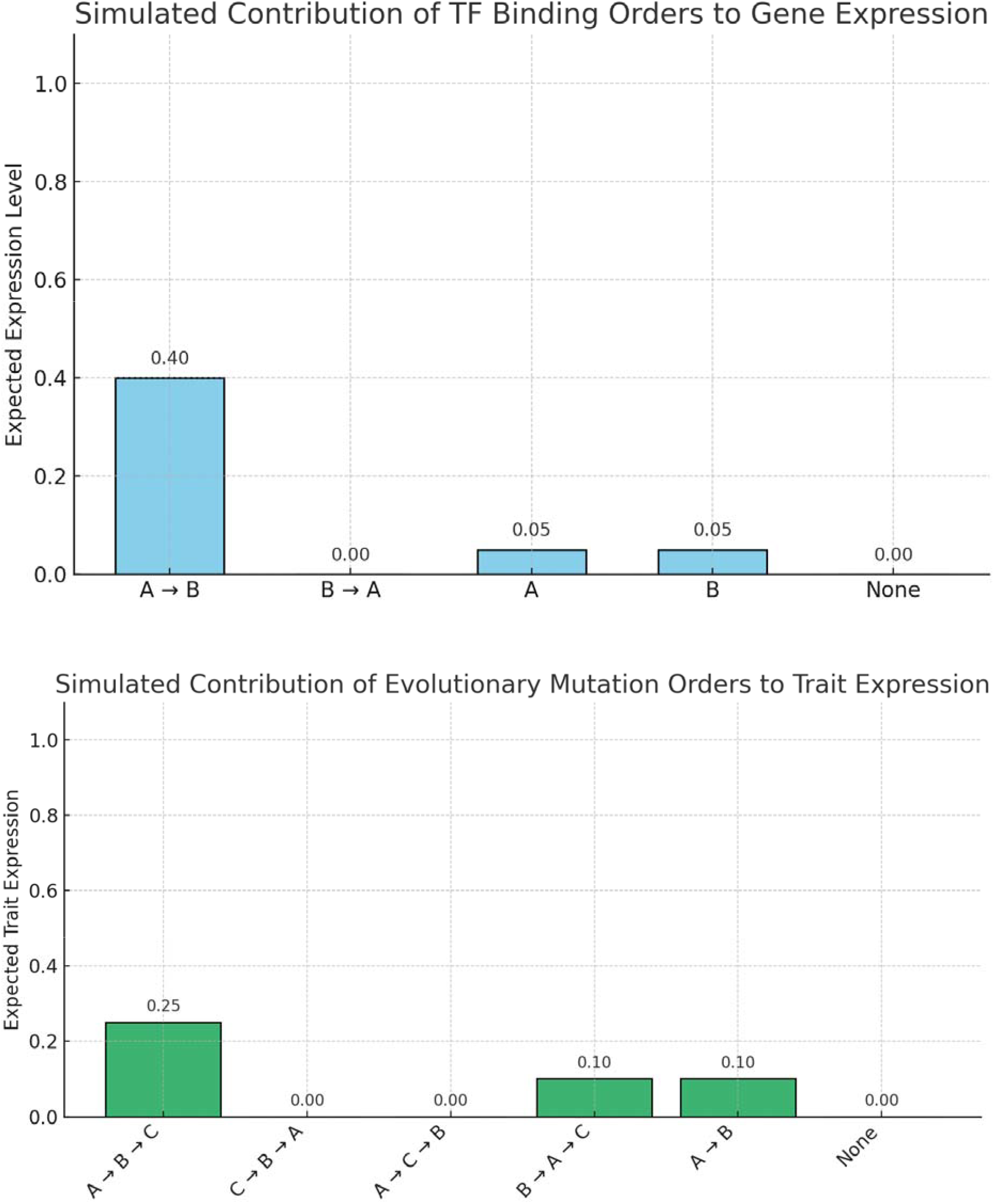
Simulation results based on algebraic modeling demonstrate the differential contributions of specific event sequences to biological and evolutionary outcomes. The two figures illustrate how variations in the order of molecular or mutational events yield distinct levels of functional expression, reflecting the non-commutative nature of the underlying processes. **Upper Figure**. Simulated plot showing the expected contribution of different transcription factor (TF) binding orders to gene expression. The x-axis represents the five binding order scenarios and the y-axis shows the expected gene expression contribution. The order A → B dominates the expression profile, while non-commutative reversals and partial bindings contribute minimally. **Lower figure**. Simulated contribution of evolutionary mutation orders to trait expression. The bar plot displays the expected trait expression resulting from six different evolutionary mutation sequences involving mutations A, B and C in different historical order. The path A → B → C is the dominant contributor to trait expression, while reversed and scrambled sequences have lower or null impacts, illustrating the importance of mutation sequence in trait emergence. Expected values were computed by weighting each outcome (ON = 1.0, PARTIAL = 0.5, OFF = 0.0) by the relative likelihood of each mutational pathway.

### Semisimplicity and radical reduction

Following the construction of the algebra 𝒜, we proceed to classify its ideal structure and determine whether it is semisimple. For this purpose, we employ the standard result from ring theory: a finite-dimensional associative algebra 𝒜 over a field 𝔽 is semisimple if and only if its Jacobson radical Rad (𝒜) is zero (Farb and Dennis, 1993; Bhuniya and Sarkar, 2023). To compute Rad (𝒜), we use the characterization of the radical as the intersection of the annihilators of all simple left 𝒜-modules. Computationally, we identify nilpotent ideals by evaluating the action of generators on test modules and examining their closure under multiplication. Letting

M be a left 𝒜-module, we compute endomorphism rings End _𝒜_ (*M*) and examine whether elements of 𝒜 act nilpotently on M, i.e., whether for each *a* ∈ 𝒜, there exists *k* ∈ ℕ such that *a*^*k*^ · *m =* 0 for all *m*∈ *M*. If Rad (𝒜) =0, then Wedderburn–Artin theory applies in full. Otherwise, we restrict to the semisimple quotient 𝒜_SS_ = 𝒜/ Rad (𝒜), which by construction is semisimple and suitable for decomposition. The establishment of semisimplicity or identification of a reduced semisimple component is critical for ensuring that Wedderburn’s theorem is applicable and yields a well-defined matrix decomposition (Behboodi et al., 2016).

### Wedderburn–Artin decomposition

Once semisimplicity is confirmed, we invoke the Wedderburn–Artin theorem, which asserts that any finite-dimensional semisimple algebra over a field 𝔽 is isomorphic to a finite direct sum of full matrix algebras over division rings (Ma et al., 2022). Explicitly, the decomposition takes the form 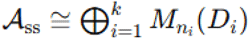, where *D*_*i*_ each is a division ring over 𝔽 and 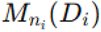 denotes the algebra of *n*_*i*_ × *n*_*i*_ matrices with entries in *D*_*i*_ i. We compute this decomposition by identifying a complete set of pairwise orthogonal primitive central idempotents {*e*_1,…,_ *e*_*k*_} ⊆ 𝒜_SS_ satisfying 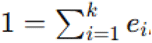 and then studying each component *e*_*i*_ 𝒜_SS_ *e*_*i*_. For each *i*, the algebra *e*_*i*_ 𝒜_SS_ *e*_*i*_ is isomorphic to 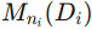 and its dimension is determined by computing the trace of the identity under the regular representation. The idempotents are constructed using polynomial identities and Peirce decomposition with computational assistance from symbolic algebra software such as GAP or SymPy (Ánh et al., 2020). This matrix block structure provides the modular decomposition of the algebra and forms the basis for associating behavioral modules with biological subfunctions.

### Representation theory and state space construction

Each matrix block 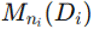 is then interpreted as an irreducible behavioral submodule of the biological system. To make this connection explicit, we construct representations ρ*i*: 𝒜_SS_ →End_*Di*_(*V*_*i*_) where is a vector space of dimension *n*_*i*_ over the division ring *D*_*i*_. These representations define the action of biological operations on state spaces and we study the structure of these actions by analyzing the module homomorphisms ϕ: 𝒜 →End(*V*). The system’s state space V is constructed based on observable outcomes (e.g., ON/OFF gene states, trait expressions, regulatory activation) and the representation maps each algebra generator to a linear operator on V. The transition structure of the system is then given by the composition of these operators, corresponding to the sequence of biological events. We explicitly compute matrix representatives for each generator using a basis adapted to the irreducible modules, enabling the translation of abstract algebraic structure into linear action on state vectors. This correspondence makes it possible to characterize each biological process as a composition of linear transformations, despite the nonlinearity of the underlying sequence logic.

### Computational implementation and algebraic tools

To implement these constructions and carry out symbolic calculations, we utilized the Python-based algebraic software SymPy (v1.12) for symbolic manipulation of non-commutative expressions, alongside GAP (Groups, Algorithms, Programming; v4.12.0) for explicit computation of group actions, idempotents and module decompositions. All matrix computations, linear maps and basis transformations were performed using NumPy and SciPy within Python (v3.11). Custom scripts were written to automate the construction of free algebras, define relation sets and compute quotient structures. The modules were constructed by evaluating the left regular representation λ: 𝒜 →End_𝔽_ (𝒜), defined by λ(*a*) (*x*) =*ax* for all *x* ∈ 𝒜. We further used this representation to identify minimal left ideals 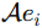, where each *e*_*i*_ is a primitive idempotent and verified the decomposition by checking that the direct sum of minimal ideals recovers 𝒜_SS_. These computational steps were necessary to ensure reproducibility of the decomposition and to precisely associate algebraic structure with observable states. With this infrastructure, we operationalized Wedderburn decomposition as a method of extracting canonical linear modules from inherently ordered and context-sensitive biological systems.

### Case studies and module-phenotype mapping

The final methodological stage involved applying this framework to case studies constructed from simplified biological systems, i.e., synthetic gene regulatory networks and abstracted evolutionary mutation paths. For each of the two cases under examination, we specified a generator set 𝒢, defined the relations R reflecting biological logic (e.g., suppression, activation or context-specific outcomes) and formed the quotient algebra 𝒜. The regular representation λ was computed explicitly and the resulting matrices were block-diagonalized according to the decomposition 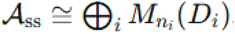. We then examined the image of each generator under the regular representation and verified its consistency with the algebraic structure. To assess whether different mutation sequences led to equivalent or distinct outcomes, we compared the orbits of state vectors under different compositions of operators. Equivalence classes of behaviors were then associated with the respective matrix blocks, allowing us to link the algebraic decomposition with functionally distinguishable system trajectories. In cases involving feedback or looped behavior, we analyzed the structure of the center *Z*(𝒜) and studied its interaction with the radical to determine whether cyclic paths contributed to reducibility or generated central idempotents. The mapping from sequence dynamics to linear modules was then systematically recorded and all operations were performed in a consistent symbolic algebraic pipeline. This last phase translated the abstract decomposition into interpretable biological substructures and allowed rigorous tracking of how different pathways were embedded in the module structure.

To simulate the effect of transcription factor (TF) binding order on gene expression (Culyba 2019; Lai et al., 2019), we implemented a probabilistic and algebraically structured model based on a simplified gene regulation scenario. This scenario represents a system where two transcription factors, A and B, interact with a gene promoter in an order-sensitive manner. The simulation aimed to quantify how different binding sequences contribute to the expected level of gene expression, based on their assigned functional outcomes and relative likelihoods.

The first step involved enumerating all meaningful binding order scenarios between TFs A and B. We defined five distinct cases: (1) A binds before B (A → B), (2) B binds before A (B → A), (3) only A binds (A), (4) only B binds (B) and (5) neither factor binds (None). These cases represent a simple but comprehensive combinatorial basis for assessing order effects in a non-commutative setting.

Next, we assigned a gene expression outcome to each binding scenario based on biological logic. In our model, the sequence A → B was considered activating, producing a full gene expression response and therefore was assigned an outcome value of 1.0. Conversely, B → A was designated as repressive or ineffective, yielding an expression value of 0.0. Scenarios in which only A or only B binds were considered partial and biologically ambiguous and each was assigned an intermediate expression value of 0.5. If neither TF binds, gene expression is assumed to remain OFF, with a value of 0.0. These scalar values serve as surrogate quantitative markers for gene activity.

To simulate how often each sequence might occur in a biological context, we introduced a discrete probability distribution over the five scenarios. These likelihoods were not derived from experimental data but chosen to reflect a plausible distribution of events in a regulated system: A → B at 40%, B → A at 30% and the remaining scenarios (only A, only B or none) each at 10%. These values sum to one and form a proper discrete probability space over which expected values can be computed.

We then computed the expected contribution of each binding scenario to overall gene expression. This was done by multiplying each outcome value by its corresponding probability, resulting in a weighted contribution.

In evolutionary biology, the order of mutations can critically shape the resulting phenotype (Platt et al., 2018; Dong et al., 2020), especially when gene interactions are epistatic. In such cases, a mutation’s effect depends on the presence or absence of earlier mutations, making the system inherently non-commutative. To simulate this, we constructed a model where a trait is determined by a sequence of mutations labeled A, B and C. The outcome of trait expression varies depending on the order in which these mutations accumulate in an evolving lineage.

We considered six distinct mutation paths: the adaptive path A → B → C which leads to a functional trait (ON), a reversed path C → B → A which yields a non-functional trait (OFF), two scrambled but plausible paths A → C → B and B → A → C (the latter leading to a partially functional trait), an incomplete path A → B and a baseline with no mutations. Each path represents a historical evolutionary sequence and the trait outcome depends not just on the presence of all mutations but on the order in which they occurred.

To simulate the impact of these pathways, we assigned a probability to each mutation sequence, reflecting its likelihood of occurring under evolutionary constraints. For instance, the most advantageous path (A → B → C) was given a higher likelihood of 0.25, while less structured or scrambled sequences received lower probabilities. Trait outcomes were modeled as 1.0 for full expression (ON), 0.5 for partial expression and 0.0 for OFF. The expected contribution of each path to the trait’s presence in a population was then calculated by multiplying the path’s probability by its trait value.

Overall, our methodological framework formalizes order-dependent biological processes within finite-dimensional non-commutative algebras, enabling their decomposition into interpretable matrix modules. Through algebraic construction, representation theory and symbolic computation, the approach yields a tractable model capturing the internal logic of complex biological and evolutionary systems with mathematical precision.

## RESULTS

The results of our simulations performed through algebraic modeling demonstrated the differential contributions of specific event sequences to biological and evolutionary outcomes, confirming the relevance of non-commutative structure in determining system behavior.

In the gene regulation model, where transcription factors A and B bind in distinct orders, the simulation showed that only the binding sequence A → B yielded a non-zero expected gene expression level of 0.40. All other permutations, including B → A and singular bindings of A or B, showed limited or no activation, with partial expression values of for each partial sequence and 0.00 for the reverse and null cases. The total expected gene expression across all scenarios, computed as the weighted sum of individual contributions, amounted to 0.50. These findings suggest that, despite the presence of multiple permissible binding sequences, a single order exerts a dominant influence on the resulting functional outcome. These differences reflect the non-commutative algebra of the transcription factor interactions and validate the use of matrix-based representation to isolate behaviorally distinct modules. Therefore, our analysis establishes a measurable structure-to-function mapping within non-commutative regulatory systems, setting the stage for evaluating more complex models.

In the evolutionary scenario, six possible mutation sequences involving A, B and C were analyzed to evaluate their relative influence on trait expression. The canonical path A → B → C contributed the highest expected value to the phenotype, at 0.25, based on a likelihood of 0.25 and a trait expression level of 1.0. The sequence B → A → C, though biologically plausible, resulted in a partial trait outcome with a weighted contribution of 0.10, while both C → B → A and A → C → B yielded zero contribution due to complete trait suppression. The incomplete path A → B, assigned a partial expression value, also contributed 0.10. Summing across all paths, the overall expected expression level for the trait was 0.45, slightly lower than that observed in the regulatory case. These results confirm that sequence order is not only a determinant of trait expression but also differentiates mutational pathways into functionally distinct classes. The algebraic decomposition of these sequences aligns with observed outcome categories, demonstrating that even in a limited path set order-specific behavior emerges with measurable frequency-dependent significance.

Overall, our results show that, in both gene regulation and evolutionary models, certain sequences dominate phenotypic expression while others contribute minimally or not at all. Algebraic structuring of operations successfully distinguished between behaviorally meaningful and redundant pathways, supporting the use of decomposition techniques in capturing the functional relevance of ordered biological processes.

## CONCLUSIONS

Our study showed that non-commutative algebraic structures may provide a rigorous framework for representing and analyzing biological and evolutionary processes that are sensitive to the order of events. In both the modeled systems, the total expected functional outcome was significantly weighted toward a single canonical sequence, confirming that decomposition captures not just theoretical structure, but also functional hierarchy. By encoding binding events and mutations as elements of a non-commutative algebra and decomposing these structures using the Wedderburn–Artin theorem, we were able to recover a modular representation of the system, where behaviorally distinct subspaces emerged naturally as matrix blocks. This approach allowed for an objective quantification of pathway contributions across multiple possible event sequences. In both the transcription factor binding model and the mutational trajectory simulation, we found that specific sequences were disproportionately responsible for functional outcomes, as demonstrated by their quantitatively dominant contributions to gene expression or trait presence. These effects were not artifacts of probabilistic assignment, but rather consequences of the algebraic formulation itself, which preserved the asymmetries introduced by ordering.

The novelty of this approach lies in the use of a formal theorem from ring theory—specifically, the Wedderburn–Artin decomposition of finite-dimensional semisimple algebras (Nakazi and Yamamoto. 2007)—as the analytic foundation for modeling non-commutative biological logic. Unlike efforts relying on probabilistic, differential or heuristic network models, our method provides a mathematically rigorous treatment of order-sensitive operations by expressing them within an associative algebra and systematically decomposing the algebra into irreducible modules. Each biological or evolutionary event is modeled as an algebraic generator and the observed or potential sequences of events form words within the free associative algebra. By imposing biologically meaningful relations and factoring the algebra accordingly, we construct a finite-dimensional object that retains the full ordering logic of the system. The application of Wedderburn’s theorem then enables the decomposition of this algebra into matrix algebras over division rings, each corresponding to a functionally irreducible substructure within the system (Kawai and Macedo Ferreira, 2024). This modularity is not imposed but emerges from the inherent algebraic structure, allowing for unambiguous classification of pathways into equivalence classes with shared functional effects. Furthermore, the use of representations allows for the direct construction of transition matrices acting on state spaces, offering a linear and tractable view of an otherwise nonlinear system (Etienne, 2015). Unlike other methods that simulate dynamics or infer networks from data, our approach works at the structural level, making it particularly suited for understanding constraint architecture, behavioral symmetries and the irreversibility of biological operations. Additionally, it provides a basis for formal comparisons between distinct systems by analyzing the isomorphism types of their corresponding algebras and modules.

Applying Wedderburn-style decomposition to biological algebras—especially those modeling non-commutative processes like gene regulation—offers practical, conceptual and computational advantages (**Figure 3**). One of the most significant advantages is the modular decomposition of complex behavior. This technique breaks down a complicated, non-commutative system into simple, well-understood components in the form of matrix blocks. This is particularly important in biology, where systems are often composed of regulatory modules (Nomiri et al., 2022). Each matrix block may correspond to a specific gene regulatory motif, signaling unit or dynamical regime such as ON, OFF or bistable states. The decomposition is analogous to isolating independent circuits in a tangled wiring diagram, which allows each part of the system to be studied and interpreted independently.

**Figure 3.**
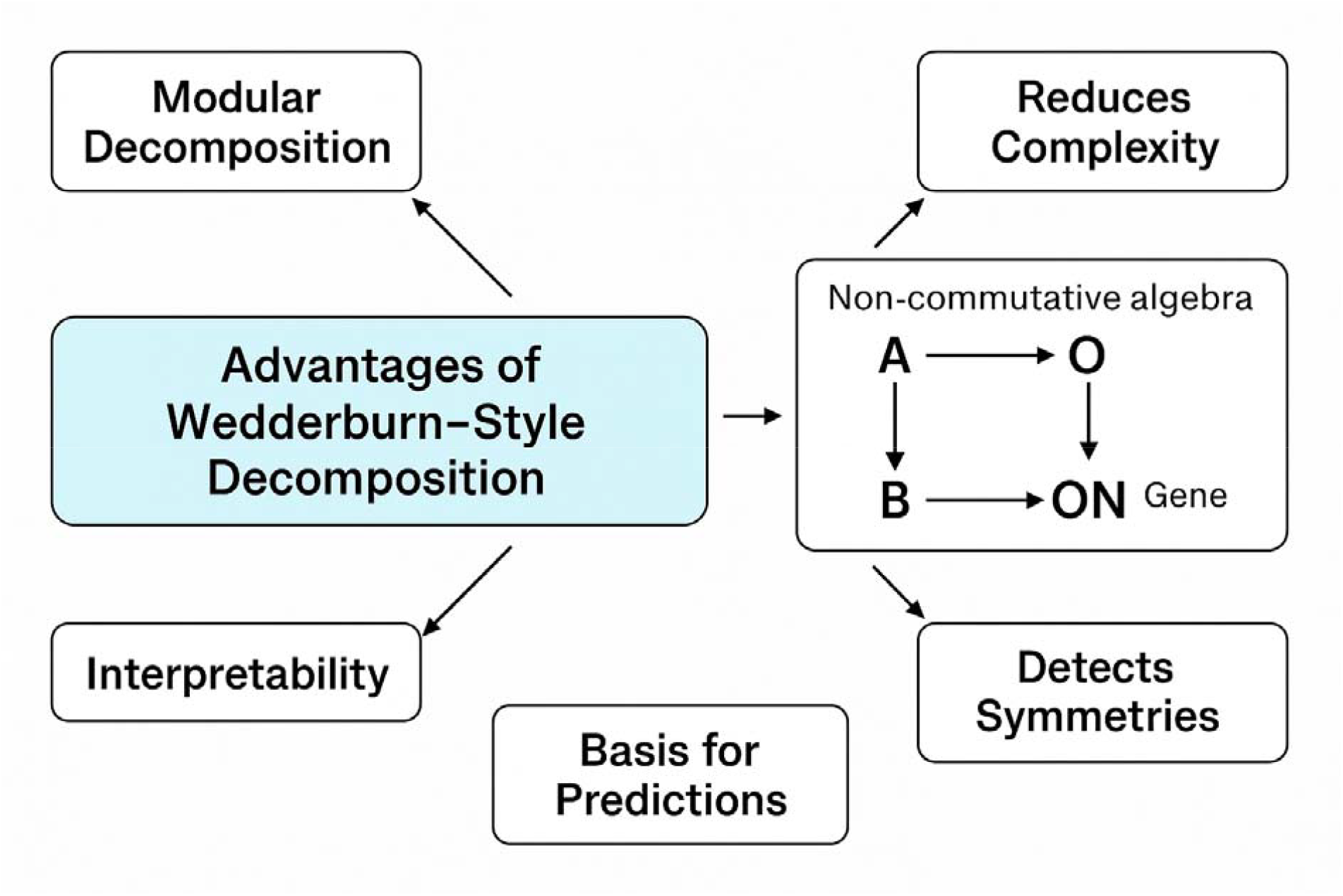
Methodological advantages of Wedderburn-style decomposition.

Another key advantage lies in the interpretability of functional units. Each irreducible component of the algebra maps to a core functional behavior. For instance, a gene being ON, OFF or in a transient or unstable expression state. This direct correspondence enables biologists to associate algebraic components with observable phenotypes, clarifying how a particular sequence of regulatory events yields specific biological outcomes. As a result, it becomes easier to interpret why certain pathways lead to activation while others do not. This level of structural insight enhances the clarity of biological modeling. Wedderburn’s decomposition also classifies all possible behaviors by providing a complete description of the system’s representations (Subroto 2024). This means that all stable, transient or observable outcomes governed by the system’s internal logic are captured and structurally distinguished.

Reducing computational complexity is another advantage. Instead of working with a large and entangled algebra, the researcher deals with smaller matrix blocks. This simplification enables faster simulation, more manageable symbolic manipulation and even more efficient parameter fitting when applied to computational biology models. This feature is particularly useful when dealing with large-scale gene regulatory networks or metabolic systems, where modular structure is a recurring theme. Internal symmetries—such as when two different sequences yield the same regulatory outcome—become clearly identifiable. This may point to underlying principles of evolutionary conservation, redundant control mechanisms or minimal sets of necessary regulatory elements. It may also suggest which transcription factors are functionally overlapping or only conditionally relevant in specific contexts.

Diverse biological systems—including neural networks, genetic interaction networks, and signal transduction pathways—that exhibit order-dependent behaviour can be analysed using a unified set of algebraic principles within a single coherent framework.

Compared to existing techniques, our algebraic framework provides advantages in terms of interpretability, modularity and structural precision. Unlike differential equation models, which rely on continuous variables and often require local linearization around equilibria (Fröhlich et al., 2019; Browning et al., 2020; Simpson et al., 2024), the present method operates in a discrete and purely structural context, preserving the full asymmetry of ordered events. Similarly, while Boolean or logical models can handle discrete transitions, they typically do not account for the algebraic consequences of sequence ordering unless explicitly coded into rules (Wang et al., 2012; Chagas et al, 2023). Probabilistic graphical models and Markov chains often collapse sequence variations into state transition probabilities, masking the importance of intermediate orderings (Mukherjee and Mitra, 2005; Pirogov et al., 2016). In contrast, our algebraic formulation explicitly tracks sequence compositions and assigns structural identities to each unique pathway. This distinction is crucial in modeling systems where path dependency determines phenotype or function.

Potential applications of this methodology span several areas of biological and evolutionary research. In genetics, the model could be used to analyze cis-regulatory modules where transcription factor order determines gene activation thresholds or combinatorial logic (Schmitz et al., 2022). By constructing algebras over known regulatory grammars and decomposing them, researchers could identify whether distinct sequences of binding events fall into equivalent outcome classes or represent unique regulatory codes. In evolutionary biology, our model could be applied to epistatic networks where mutations exhibit conditional effects depending on genetic background (Anholt 2020; Lin, 2021). In these cases, evolutionary trajectories could be encoded algebraically and decompositions could reveal redundant or convergent paths. Moreover, in developmental biology, cell differentiation programs could be abstracted as sequences of transcriptional decisions and the decomposition would allow identification of minimal irreducible trajectories required for cell fate specification. From an experimental standpoint, the model supports testable hypotheses: for example, perturbing a specific binding order predicted to be functionally critical (i.e., corresponding to a distinct irreducible module) should result in a measurable phenotypic shift, while perturbing sequences within a common block should not. Understanding the algebraic structure allows for predictions such as what might happen when a specific transcription factor is knocked out, which sequence orders are crucial for gene activation and whether any intermediate or partial activation states exist within the system. These predictions may be especially relevant for experimental design in systems and synthetic biology.

Furthermore, constructing the full algebra from empirical or inferred interaction networks and then applying Wedderburn decomposition could enable experimentalists to test whether observed biological behavior aligns with predicted structural modules. This opens the possibility of designing experiments to distinguish between systems that are functionally robust (i.e., share algebraic decompositions) versus those that are structurally fragile.

Several limitations should be acknowledged. Our framework requires complete specification of the generator set and the relations governing the algebra, which may not always be fully known or measurable in biological contexts. Still, the method is designed for finite-dimensional algebras and may not directly generalize to continuous systems or those requiring infinite state representations. While certain extensions may be possible through projective limits or infinite-dimensional representations, these go beyond the scope of this study. Althoughour approach captures order sensitivity, it does not inherently model timing, rates or dynamics unless these are separately encoded into state transitions. Thus, our method complements but does not replace dynamical systems theory or probabilistic inference when continuous modeling is essential. Additionally, computing the Wedderburn decomposition symbolically for large algebras may be computationally intensive, particularly when the algebra has many generators or complex non-trivial relations. While our approach is powerful for mid-scale systems, it may require additional simplification strategies for larger biological networks. A final consideration is the interpretability of division rings in biological terms, especially when representations involve fields other than ℝ or C. The mathematical coherence remains intact, but the biological semantics of such structures may not be immediately clear.

In conclusion, we demonstrate that finite-dimensional non-commutative algebras and Wedderburn decomposition offer a mathematically sound framework for modeling biological systems in which the order of operations is functionally significant.

## DECLARATIONS

### Ethics approval and consent to participate

This research does not contain any studies with human participants or animals performed by the Author.

### Consent for publication

The Author transfers all copyright ownership, in the event the work is published. The undersigned author warrants that the article is original, does not infringe on any copyright or other proprietary right of any third part, is not under consideration by another journal and has not been previously published.

### Availability of data and materials

All data and materials generated or analyzed during this study are included in the manuscript. The Author had full access to all the data in the study and took responsibility for the integrity of the data and the accuracy of the data analysis.

## Acknowledgements

None.

## Authors’ contributions

The Author performed: study concept and design, acquisition of data, analysis and interpretation of data, drafting of the manuscript, critical revision of the manuscript for important intellectual content, statistical analysis, obtained funding, administrative, technical and material support, study supervision.

## Declaration of generative AI and AI-assisted technologies in the writing process

During the preparation of this work, the author used ChatGPT 4o to assist with data analysis and manuscript drafting and to improve spelling, grammar and general editing. After using this tool, the author reviewed and edited the content as needed, taking full responsibility for the content of the publication.

